# Downregulation of ribosomal RNA (rRNA) genes in human head and neck squamous cell carcinoma (HNSCC) cells is linked to rDNA promoter hypermethylation

**DOI:** 10.1101/2023.03.13.532507

**Authors:** Neha Priyadarshini, Navinchandra Venkatarama Puppala, Jayasree Peroth Jayaprakash, Piyush Khandelia, Vivek Sharma, Gireesha Mohannath

## Abstract

Eukaryotes carry hundreds of ribosomal RNA (rRNA) genes as tandem arrays, which generate rRNA for protein synthesis. Humans carry ~ 400 rRNA gene copies, which are epigenetically regulated. Dysregulation of rRNA synthesis and ribosome biogenesis are characteristic features of cancers. Targeting aberrant rRNA expression for cancer therapy is being explored. Head and neck squamous cell carcinoma (HNSCC) is among the most prevalent cancers globally. Using quantitative PCR and bisulfite sequencing, we show that rRNA genes are downregulated and their promoters are hypermethylated in HNSCC cell lines. These finding may have relevance for prognosis and diagnosis for HNSCC.

## Introduction

The ribosomes are the sites of cellular protein synthesis, and the ribosomal RNA (rRNA) synthesis rate is tightly linked to cellular growth and proliferation. In eukaryotes, ribosomes are made up of four catalytic rRNAs and ~ 80 protein subunits [1]. Eukaryotic genomes encode hundreds to thousands of rRNA genes that occur as tandem arrays at chromosomal locations called nucleolus organizer regions (NORs) [2–5]. Nucleoli are rRNA transcription and processing sites and ribosomal subunits are assembled here [6–10]. In humans, ~ 400 copies of 45S rRNA genes are distributed among five NORs that are located one each on acrocentric chromosomes 13, 14, 15, 21, and 22 (Fig. 1) [11]. Each human 45S rRNA gene is ~ 43 kb, including ~ 13 kb pre-45S rRNA encoding sequences (5’ ETS, 18S, ITS1, 5.8S, ITS2, 28S, and 3’ ETS) preceded by a promoter and a ~ 30 kb intergenic spacer [12]. RNA polymerase I (RNA Pol I) transcribes 45S rRNA genes to produce pre-45S rRNA, also known as 47S rRNA (throughout this manuscript, we referred to it as 45S rRNA), which is then processed to 18S, 5.8S, and 28S rRNAs, which form the catalytic cores of ribosomes (Fig. 1)[13, 14]. The fourth rRNA, 5S, is encoded by 5S rRNA genes which are transcribed by RNA polymerase III (Pol III), and in human cells, they occur as tandem repeats on chromosome 1 (Fig. 1) [15, 16].

**Figure 1:**
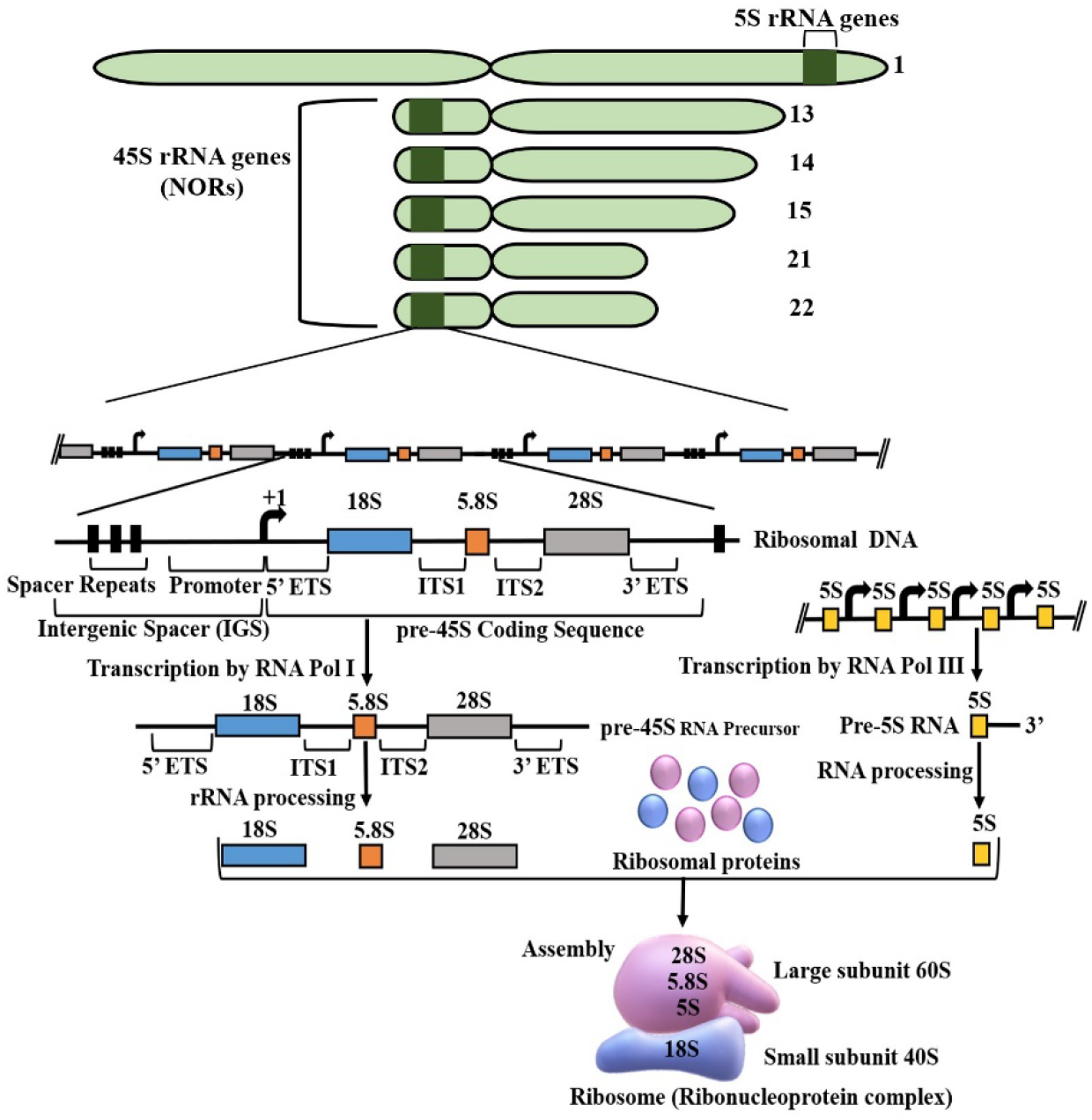
Organization of human Nucleolus Organizer Regions (NORs). The top part of the diagram shows the location of NORs containing 45 S rRNA genes on five acrocentric chromosomes (chromosome# 13, 14, 15, 21, and 22) and the locus containing 5S rRNA genes on chromosome 1. The bottom part of the diagram shows a representative tandem array of 45 S and 5S rRNA genes, the composition of an individual 45 S rDNA unit, transcription and processing of 45 S and 5S rRNA genes, and the assembly of ribosomal subunits to form mature ribosomes.

The balance between cell growth and ribosome production is maintained in normal cells by transcriptional regulation of rRNA genes at an appropriate level, either by coarse control that is accomplished via epigenetic mechanisms, such as DNA methylation and histone modifications, modulate the number of rRNA genes that are expressed [17–21] or by fine-tuning rRNA transcription rate via regulating the number of RNA Pol I enzymes engaged in transcription at each active gene [11, 22]. Recent studies in plants have shown that rRNA dosage control predominantly occurs at the level of NORs, not at the individual gene level [23, 24]. Although NOR-level regulation of rRNA genes could occur in other eukaryotes, including humans, it remains to be investigated [25].

In human cancer cells, the balance between cell growth and ribosome production is perturbed, and rRNA synthesis is dysregulated [26–28]. In multiple cancers, rRNA upregulation and/or rDNA promoter hypomethylation have been observed [29–34]. Rapidly proliferating cancer cells have enhanced metabolism with a need for efficient protein synthesis, and increased rRNA synthesis likely aids in increased ribosome biogenesis leading to a higher rate of protein synthesis. Therefore, RNA Pol I transcription is targeted as an anticancer modality in these cancers [35–46].

Interestingly, rRNA downregulation and/or rDNA promoter hypermethylation have also been observed in some cancers [47–53]. The rRNA dysregulation and tumorigenesis are not always been linked to altered rDNA promoter methylation status [32, 48, 49, 53]. Overall, several studies indicate that the effects of carcinogenesis and cellular response to carcinogenesis on rRNA gene expression in different cancers is not well-studied. Therefore, it is essential to thoroughly characterize the expression patterns of rRNA genes and the epigenetic status of their promoters in each cancer type.

HNSCC comprises malignancies of the upper aerodigestive tract, including paranasal sinuses, nasopharynx, oropharynx, hypopharynx, larynx, oral cavity, and nostrils [54–56]. Globally, it is one of the most prevalent cancers, with an estimated incidence of around 9,30,000 new cases and 4,60,000 deaths in 2020 [57]. In this study, we analyzed expression levels of rRNA genes and DNA methylation status of rDNA promoters in SCC-25, CAL-27, and FaDu HNSCC cell lines. Het-1A was used as a control cell line in this study. Our results indicate that rRNA genes are downregulated in the HNSCC cell lines due to rDNA promoter hypermethylation.

## Material and methods

### Cell culture

The human normal squamous epithelial cell line Het-1A and HNSCC cell lines SCC-25 (ATCC CRL-1628), CAL-27 (ATCC CRL-2095), FaDu (ATCC HTB-43), and a control cell line Het-1A (ATCC CRL-2692) were purchased from American Type Culture Collection (Manassas, VA, USA). Het-1A cells were cultured in LHC-9, SCC-25 in DMEM-F12, CAL-27 in DMEM, and FaDu in EMEM media (ThermoFisher Scientific) at 37°C in a humidified incubator with 5% CO2. All the HNSCC cell media were supplemented with 10% fetal bovine serum (FBS, Gibco), 1% Penicillin (Gibco), and 1% streptomycin (Gibco). The medium was changed every 2-3 days, and the cells were subcultured before a confluent monolayer could be formed. For detaching the cells, 0.25% Trypsin-EDTA was used. Cells grown to 70-80% confluence were used for isolating total RNA and DNA. Het-1A cells were grown in cell culture dishes precoated with a mixture of collagen (0.03 mg/ml), fibronectin (0.01 mg/ml), and bovine serum albumin (0.01 mg/ml) dissolved in a culture medium.

### RNA isolation and quantitative PCR

Total RNA was isolated from all the cell lines using AllPrep DNA/RNA/Protein Mini Kit (Qiagen). The total RNA was then treated with Turbo DNA-free kit (Invitrogen) to eliminate contaminating genomic DNA. DNA-free RNA was reverse transcribed to cDNA using SuperScript III First Strand cDNA synthesis kit (Invitrogen). Before performing qPCR assays, standard curve analysis was carried out to ensure that the amplification efficiency of PCR for each primer pair was within the prescribed range of 90-110% (Applied Biosystems). Quantitative PCR (qPCR) was performed using a Step One Plus real-time PCR system (Applied Biosystems) in the presence of TB Green^®^ Premix Ex Taq™ (Tli RNase H Plus) (Takara). qPCR amplification was carried out using an initial denaturation at 95°C for 10 sec, followed by 40 cycles of 15 sec at 95°C and 60 sec at 60°C. The relative gene expression of precursor 45S RNA, and mature 18S, 5.8S, 28S, 5S RNA and U6 snRNA was calculated using the 2^(-ΔΔCT)^ method [58] and the expression levels were normalised to the expression of glyceraldehyde-3-phosphate dehydrogenase (GAPDH) as a reference gene. qPCR was performed in triplicates for each sample. Primer sequences used to amplify pre-45S RNA, mature 18S, 5.8S, and 28S rRNAs are the same as those used in a previous study by Uemura et al. [32]. The primers used in all the qPCR reactions are listed in Figure S1.

### Bisulfite sequencing of the rDNA promoter

DNA was isolated from all the cell lines using AllPrep DNA/RNA/Protein Mini Kit (Qiagen). The samples were treated with sodium bisulfite to convert unmethylated cytosine residues to uracil via deamination using the EpiTect Bisulfite kit (Qiagen). Using bisulfite-treated DNA as template, PCR amplification of the rDNA promoter region for each cell line was carried out with initial denaturation for 4 min at 95°C, followed by 35 cycles of 30 sec at 94°C, 45 sec at 45°C, and 45 sec at 72°C, and a final extension for 10 min at 72°C, in the presence of GoTaq^®^ Green Master Mix (Promega). For untreated DNA (as controls), PCR amplification was carried out with an initial denaturation for 3 min at 95°C, followed by 35 cycles of 30 sec at 95°C, 30 sec at 60°C, and 30 sec at 72°C, and a final extension for 3 min at 72°C, in the presence of GoTaq^®^ Green Master Mix (Promega). In both cases, each PCR reaction consisted of an initial denaturation for 3 or 4 min at 94°C or 95°C and a final extension of 3 or 10 min at 72°C. The PCR primers used to amplify the DNA promoter region with cytosine-to-uracil (thymine) conversions were the same as previously published [34]. The PCR products were purified using QIAquick PCR and Gel Cleanup Kit (Qiagen) and cloned into a pGEM-T Easy Vector System 1 (Promega). Plasmid DNA from ~ 20 positive clones for each cell line was isolated using QIAprep Spin Miniprep Kit (Qiagen) and the cloned rDNA promoter region was sequenced using Sanger sequencing. To ensure accuracy of the sequencing data, each clone was sequenced from both ends of the cloned promoter (forward and reverse sequencing). The methylation status of cytosines in CG, CHG and CHH contexts for each position (vertical comparison) and across the rDNA promoter (horizontal comparison) was tabulated based on the cytosine-to-uracil (thymine) conversion data. The tabulated data were then used for generating graphs. The primers used to amplify the rDNA promoter region are listed in Figure S1.

### Statistical analysis

Data from three independent biological replicates were used for all the calculations and to generate error bars. The standard t-test was used to analyze significant differences in gene expression levels and rDNA promoter cytosine methylation levels between the control and the HNSCC cell lines. A P of < 0.05 was considered statistically significant.

## Results

### The rRNA genes are downregulated in human HNSCC cell lines

To assess the expression levels of rRNA, total RNA was isolated from the HNSCC and control cell lines, and subjected to qPCR analysis. Results of qPCR assays showed a statistically significant reduction in the levels of pre-45S RNA and 18S, 5.8S, and 28S mature rRNAs. The 45S precursor is very short-lived and therefore, levels of pre-45S rRNA are considered as an indication of 45S rDNA transcription. Accordingly, our data indicate reduced 45S rDNA transcription and the steady-state levels of the mature rRNA in human HNSCC lines compared to the control cell line (Fig. 2, A-D). The reduction in rRNAs among the three HNSCC lines varied from 60.1-92.1% (pre-45S rRNA), 20.2-47.5% (18S), 47.5-77.8% (5.8S), 14.3-24.4% (28S). Further, we analyzed expression levels of 5S rRNA and U6 snRNA. We saw similar downregulation of these genes in the HNSCC cell lines (except for U6 snRNA in SCC-25 line) compared to the control (Fig. 2, E-F). The 5S rRNA and U6 snRNA reduction among the three HNSCC lines varied from 42.0-71.8 % and 31.8-48.1%, respectively. Our data show downregulation of RNA Pol I and RNA Pol III-transcribed rRNAs in the human HNSCC cell lines.

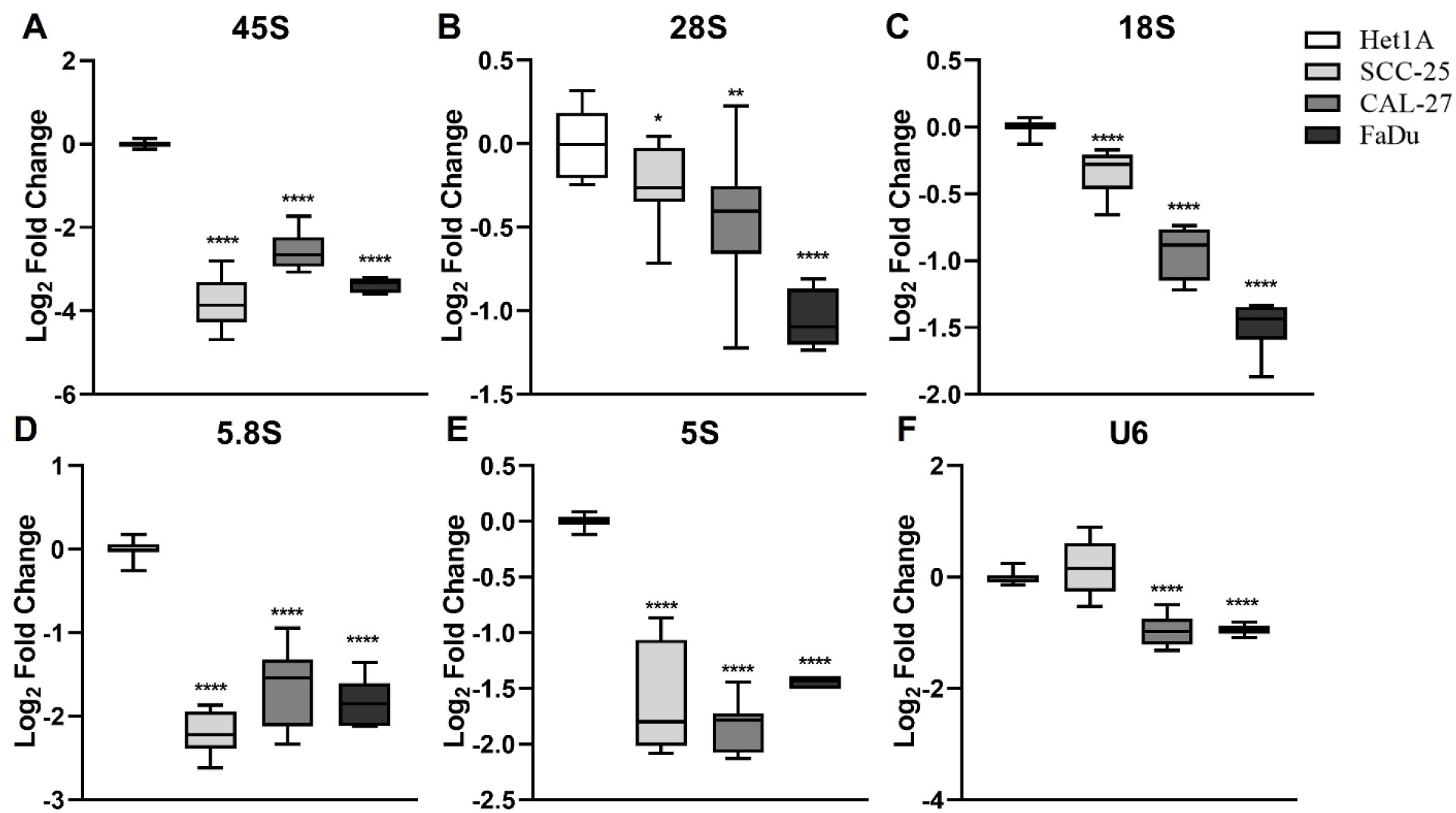

### 45S rRNA downregulation in human HNSCC cell lines correlates with rDNA promoter hypermethylation

To determine if the reduction in 45S rRNA levels in HNSCC cell lines is linked to rDNA promoter methylation (cytosine methylation), we performed sodium bisulfite sequencing of the 45S rRNA gene promoter. For this analysis, DNA isolated from the HNSCC, and the control cell lines was treated with sodium bisulfite, followed by PCR amplification and cloning of 45S rDNA promoter. Sequencing data of the rDNA promoter from the untreated DNA samples revealed that the sequence is 100% identical to the previously published sequence (Fig. 3A) [32, 34].

**Figure 3.**
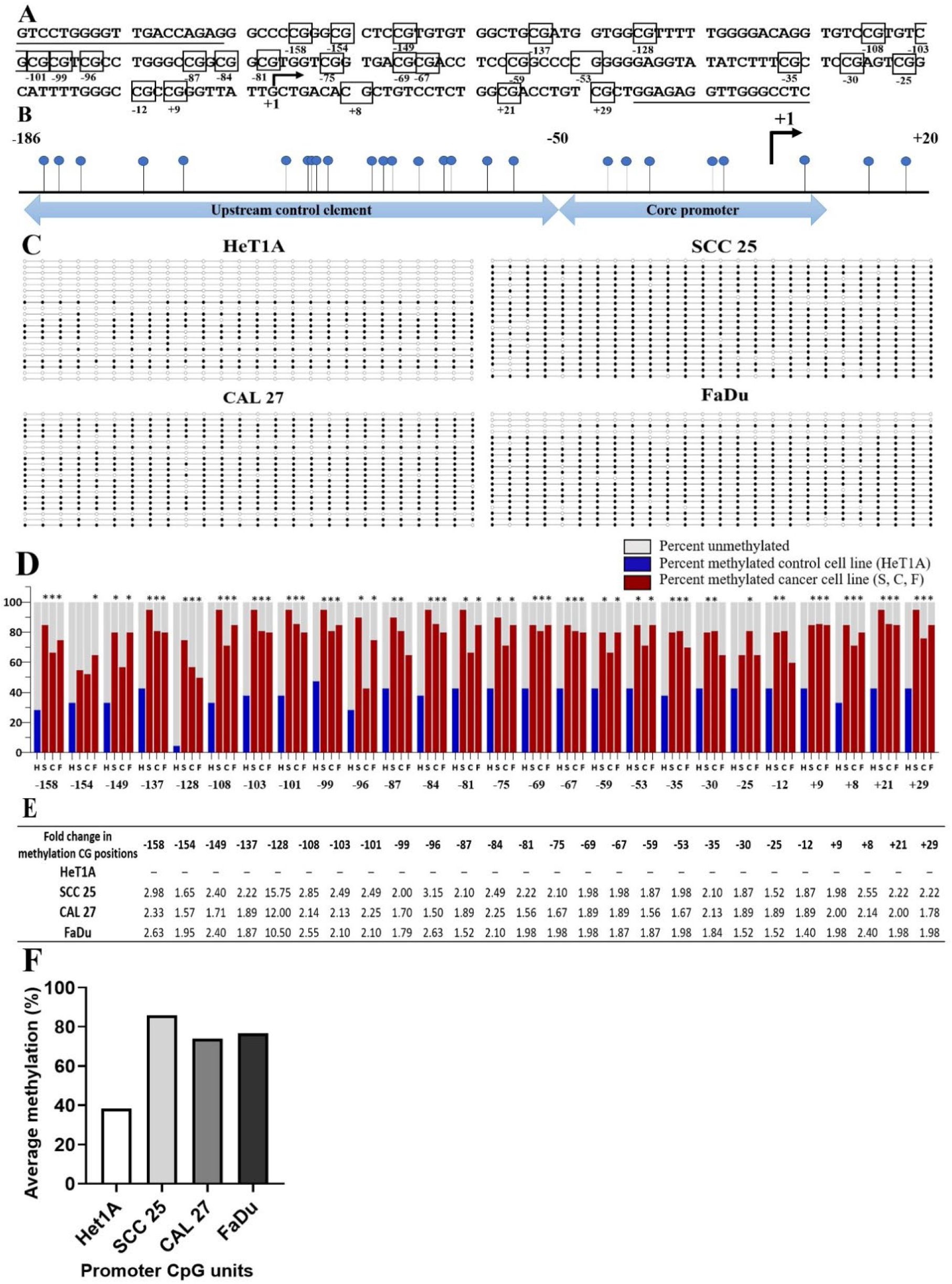
Sodium bisulfite sequencing revealed the hypermethylation of rDNA promoters in the HNSCC cell lines (SCC-25, CAL-27, and FaDu) compared to the control cell line (Het1A). (A) The rDNA promoter sequence obtained by Sanger sequencing of clones of untreated DNA of the HNSCC cell lines, the control cell line (HeT1A) and human blood. The underlined sequences represent primer binding sites. The boxes indicate the location of cytosine in CG context, in reference to the transcription start site indicated by an arrow and ‘+1’. (B) Each lollipop illustrates the position of CG dinucleotides in relation to the transcription start site (indicated by an arrow and ‘+1’) on the 45S rDNA promoter. (C) A diagram showing the methylation status of each cytosine in a CG context, with each row denoting one clone. An open circle represents unmethylated cytosines whereas a full circle represents methylated cytosines. (D) A bar diagram showing methylation density for each cytosine in a CG context for HNSCC and the control cell line (HeT1A). (E) A table showing a fold change in methylation densities for each cytosine in a CG context in HNSCC lines compared to the control cell line. (F) Average rDNA promoter methylation percentage in HNSCC cell lines and the control cell line. * indicates P<0.05.

In mammals, cytosine is predominantly methylated in CG context (also called CpG) unlike plants, where cytosine is methylated in all three contexts (CG, CHG, CHH, where H denotes any nucleotide other than G) [18]. The rDNA promoter has a total of 26 cytosines in a CG context, while the other cytosines are either in a CHG or a CHH context (Fig. 3A). Relative to the transcription start site of a 45S rRNA gene, the locations of 26 cytosines in a CG context, the core promoter sequence, and the upstream regulatory element are indicated in Figure 3B [34]. The bisulfite sequencing data revealed that most of the 26 cytosines in CG context (vertical comparison) in the HNSCC cell lines are hypermethylated compared to the control cell line with a statistical significance (Fig. 3 C-F). Most of the cytosine residues in the CG context showed more than a two-fold increase in methylation (Fig. 3E). Similarly, the HNSCC lines showed a higher percentage of rDNA hypermethylation when cytosines in the CG context were considered across the rDNA promoter (horizontal comparison) (Fig. 3F). However, we found either no methylation or very less methylation of cytosines in CHG and CHH contexts (Fig. S3). Collectively, these data indicate that the downregulation of rRNA genes in the HNSCC lines is likely mediated by rDNA promoter hypermethylation (CG methylation).

## Discussion

Ribosomal RNAs serve as the catalytic ribozyme component of ribosomes in protein biosynthesis [59]. The rRNA transcription is tightly regulated according to the cellular demands for protein synthesis. Accordingly, rRNA levels are regulated either at the level of rRNA gene dosage via epigenetic regulatory mechanisms or at the level of rDNA transcriptional rates, or both [11, 17–22]. rDNA transcription and ribosome biogenesis are dysregulated in cancer [26–28]. In some cancers, such as cervical, prostate, breast, colon, and alveolar rhabdomyosarcoma, rRNAs are upregulated with or without the corresponding rDNA promoter hypomethylation [29–34]. It is believed that rapidly growing cancer cells would require elevated protein synthesis and hence a corresponding increase in the rRNA levels. Therefore, inhibition of RNA Pol I transcription is being explored as an anticancer therapy [35–46]. However, such a modality may not be suitable for all cancers as rRNA downregulation has also been witnessed in some cancers, including breast, oral, ovarian, endometrial, and myelodysplastic syndrome [47–53]. Our study makes to this list of cancers where rRNA downregulation and the corresponding rDNA hypermethylation have been observed.

In all the HNSCC lines, we observed significant downregulation of rRNAs that are transcribed by RNA Pol I (45S rRNA) and Pol III (5S rRNA) (Fig. 2). Our qPCR-based expression analysis included both pre-45S (an indication of 45S rRNA transcription), mature rRNAs (5S, 5.8S,18S, and 28S) and U6 snRNA (another RNA Pol III target). Moreover, sodium bisulfite sequencing data revealed significant rDNA promoter hypermethylation in all three HNSCC lines compared to the control cell line (Fig. 3), which is likely the mechanism underlying the observed rRNA downregulation in these lines. Cytosine methylation was mostly found in CG context (Fig. 3C-F) and either no or to a negligible extent in CHG and CHH contexts (Fig. S2). This is in contrast to findings from a study on oral squamous cell carcinoma (OSCC), where downregulation of rRNA has been reported with unchanged rDNA promoter methylation [48]. One potential reason for not seeing any change in rDNA promoter methylation in this study, as speculated by the authors, is that in these patients with regular intake of tobacco and alcohol, histologically classified normal cells might have undergone epigenetic alterations in the tissues surrounding the tumor, and these cells might have a potential malignant transformation but have not yet developed a fully neoplastic phenotype [48]. Alternatively, the sample size could have been smaller (12 samples) [48]. Large sample size in these studies could be critical as signified by a breast cancer study involving 68 samples where in some patients showed a correlation between enlarged nucleolar size (an indication of enhanced rRNA expression) and rDNA promoter hypermethylation while a subgroup of patients did not show such a correlation [49]. Moreover, another study on breast cancer reported rather an upregulation of rRNAs [30]. In light of these reports, our findings highlight the need to carry out more studies on HNSCC patients with a larger sample size to better understand the link between rRNA downregulation and carcinogenesis.

One important question remains to be answered; why are rRNA levels reduced in some cancers? A convincing answer is yet to emerge, but we speculate two possible reasons. In some cancers, formation of specialized ribosomes termed ‘onco-ribosomes’ may drive preferential translation of selective mRNA that would favor the proliferation of the cancer cells [60–62]. In such a case scenario, limited expressed rRNA would be sufficient to translate oncogenes that are necessary for cell proliferation rather than upregulating ribosomal RNA to synthesize all proteins. Secondly, the downregulation of rRNA could result from a host response to cause apoptosis either in a p53-dependent [39, 63–65] or a p53-independent manner [66]. However, further experimentation is needed to test these hypotheses.

Future studies involving a comprehensive analysis of rRNA levels and rDNA promoter methylation status each cancer type with large sample size are warranted. Such studies can potentially develop rRNA level assays as cancer-specific prognostic and diagnostic markers for early detection and treatment, in addition to enabling therapeutic targeting of rRNA synthesis as an anticancer tool.

## List of abbreviations

rRNA: Ribosomal RNA
rDNA: Ribosomal DNA
ITS: Internal transcribed spacers
ETS: External transcribed spacers
HNSCC: Head and neck squamous cell carcinoma
NOR: Nucleolus organizer regions
snRNA: Small nuclear RNA
GAPDH: Glyceraldehyde-3-phosphate dehydrogenase
CG: Cytosine Guanine
CHG: Cytosine (H is any nucleotide other than guanine) Guanine
CHH: Cytosine (H is any nucleotide other than guanine)

## Author contributions

N.P. N.V.P., P.K., V.S., and G.M. designed research, N.P., N.V.P., and J.P.J carried out research, N.P., N.V.P., P.K., V.S. and G.M. analyzed data, N.P., N.V.P., and G.M. wrote the manuscript. All authors reviewed the manuscript.

## Acknowledgements

The authors are also thankful to all those from the biological sciences and other departments at BITS-Pilani Hyderabad campus, who have supported this work in one way or another.

