## Supplementary figures for "Downregulation of ribosomal RNA (rRNA) genes in human head and neck squamous cell carcinoma (HNSCC) cells is linked to rDNA promoter hypermethylation"

| *Primer* | *Forward (5'-3')* | *Reverse (5'-3')* |
|---|---|---|
| 47S rRNA | GAACGGTGGTGTGTCGTT | GCGTCTCGTCTCGTCTCACT |
| 28S rRNA | AGAGGTAAACGGGTGGGGTC | GGGGTCGGGAGGAACGG |
| 18S rRNA | GATGGTAGTCGCCGTGCC | GCCTGCTGCCTTCCTTGG |
| 5.8S rRNA | ACTCGGCTCGTGCGTC | GCGACGCTCAGACAGG |
| 5S rRNA | CCATACCACCCTGAACGC | AGCACCCGGTATTCCCAG |
| U6 snRNA | CTCGCTTCGGCAGCACA | AACGCTTCACGAATTTGCGT |
| GAPDH | CGCTCTCTGCTCCTCCTGTT | CCATGGTGTCTGAGCGATGT |
| rDNA promoter (Bisulfite treated) | GTTTTTGGGTTGATTAGA | AAAACCCAACCTCTCC |
| rDNA promoter (Bisulfite untreated) | CCGAAAATGCTTCCGGCTC | GCGAGAGAACAGCAGGC |

**Figure S1:** List of primers used in this study

**A**

| | **CHG-Position methylation (%)** | | | | | | | | | | | | | | | | |
|---|---|---|---|---|---|---|---|---|---|---|---|---|---|---|---|---|---|
| | **-159** | **-150** | **-140** | **-116** | **-109** | **-93** | **-88** | **-60** | **-54** | **-31** | **-13** | **-10** | **+2** | **+10** | **+17** | **+25** | **+31** |
| **HeT1A** | 0 | 0 | 0 | 0 | 0 | 0 | 0 | 0 | 0 | 0 | 0 | 0 | 0 | 0 | 0 | 0 | 14 |
| **SCC 25** | 0 | 10 | 15 | 0 | 0 | 9.5 | 0 | 5 | 10 | 5 | 0 | 0 | 5 | 10 | 10 | 0 | 10 |
| **CAL 27** | 4.8 | 0 | 4.8 | 0 | 0 | 0 | 0 | 0 | 0 | 0 | 0 | 0 | 4.8 | 4.8 | 4.8 | 0 | 9.5 |
| **FaDu** | 15 | 0 | 0 | 15 | 0 | 15 | 0 | 0 | 5 | 0 | 0 | 0 | 0 | 5 | 5 | 0 | 0 |

**B**

| | **Average promoter methylation (%)** |
|---|---|
| **HeT1A** | 0.89 |
| **SCC 25** | 5.62 |
| **CAL 27** | 2.08 |
| **FaDu** | 3.75 |

**C**

| | **CHH-Position methylation (%)** | | | | | | | | | | | | | | | |
|---|---|---|---|---|---|---|---|---|---|---|---|---|---|---|---|---|
| | **-161** | **-160** | **-152** | **-94** | **-64** | **-63** | **-61** | **-56** | **-55** | **-39** | **-33** | **-22** | **+6** | **+14** | **+15** | **+24** |
| **HeT1A** | 0 | 0 | 4.8 | 0 | 0 | 0 | 0 | 0 | 0 | 0 | 0 | 0 | 0 | 0 | 0 | 0 |
| **SCC 25** | 5 | 5 | 0 | 5 | 0 | 10 | 0 | 5 | 0 | 15 | 0 | 0 | 0 | 5 | 0 | 5 |
| **CAL 27** | 0 | 0 | 0 | 4.8 | 0 | 4.8 | 0 | 0 | 0 | 0 | 0 | 0 | 0 | 0 | 0 | 0 |
| **FaDu** | 0 | 0 | 0 | 0 | 0 | 0 | 0 | 0 | 0 | 0 | 0 | 0 | 0 | 0 | 0 | 0 |

**D**

| | **Average promoter methylation (%)** |
|---|---|
| **HeT1A** | 0.89 |
| **SCC 25** | 3.43 |
| **CAL 27** | 1.19047619 |
| **FaDu** | 1.875 |

**Figure S2:** (A) The percentage of methylation at each cytosine position in a CHG context. (B) Average rDNA promoter cytosine methylation (in CHG context) percentage for each cell line. (C) The percentage of methylation at each cytosine position in a CHH context. (D) Average rDNA promoter cytosine methylation (in CHH context) percentage for each cell line.
